# Linking the Resistome and Plasmidome to the Microbiome

**DOI:** 10.1101/484725

**Authors:** Thibault Stalder, Max Press, Shawn Sullivan, Ivan Liachko, Eva M. Top

## Abstract

The rapid spread of antibiotic resistance is a serious human health threat. A range of environments have been identified as reservoirs of the antibiotic resistance genes (ARGs) found in pathogens. However, we lack understanding of the origins of these ARGs and their spread from environment to clinic. This is partly due to our inability to identify the bacterial hosts of ARGs and the mobile genetic elements that mediate this spread, such as plasmids and integrons. Here we demonstrated that the *in vivo* proximity ligation method Hi-C can determine the *in situ* host range of ARGs, plasmids, and integrons in a wastewater sample by physically linking them to their host chromosomes. Hi-C detected both previously known and novel associations between ARGs, mobile elements and host genomes, mostly validating this method. A better identification of the natural carriers of ARGs will aid the development of strategies to limit resistance spread to pathogens.

## Main body of text

Multi-drug resistant pathogens are increasing in prevalence worldwide [1–3]. The alarming rate at which bacteria adapt to antimicrobial compounds is partly due to their ability to acquire antimicrobial resistance genes (ARGs) through horizontal transfer of mobile genetic elements such as plasmids. For example, plasmid-mediated resistance has emerged against quinolones [4], carbapenems [5], and colistin [6]. In addition, other genetic elements such as integrons facilitate the acquisition and expression of ARGs [7].

Numerous studies have revealed the diversity, abundance, and distribution of ARGs in habitats such as soil, rivers, human and animal guts, and wastewater treatment plants, implicating them all as plausible reservoirs for ARGs [8]. While cultivation-independent metagenomics of environmental samples has become a widespread approach, most metagenomic data do not provide information about the bacterial hosts of the ARGs and the replicons that carry them (chromosome vs. plasmid) [9]. They cannot identify the hosts of plasmids because total DNA extraction disconnects mobile elements from their host genomes. This host-plasmid association is nevertheless critical to understand the ecology of antibiotic resistance and the trajectories that bring resistance genes into the clinic [10].

Proximity-ligation methods such as Hi-C and 3C have been used to detect interactions between DNA molecules originating in the same cell within microbial communities (Fig. S1). Both methods are able to reconstruct strain- and species-level genomes from mixed bacterial cultures, and correctly link plasmids and phage to their bacterial hosts [11–13]. These methods can also reconstruct metagenome-assembled genomes (MAGs) from bacterial communities such as those of the mammalian gut communities [14–16]. In this study, we showed that cultivation-independent metagenomic Hi-C data can help determine the reservoirs of ARGs and the plasmids and integrons that carry them in a diverse wastewater community. It also led to the reconstruction of several novel MAGs.

### Proximity ligation reconstructs a known plasmid-host association from a wastewater sample

We first validated the ability of Hi-C to assemble the genome of a completely sequenced plasmid-bearing bacterium from a wastewater metagenome. We added ~7 × 10^7^ CFU/mL of *E. coli* K12::gfp containing the multi-drug resistance plasmid pB10::rfp, hereafter named EC, to a portion of a wastewater sample (the raw sample is designated WW, the spiked sample WWEC). For both samples we generated short-read metagenome assemblies and used ProxiMeta Hi-C deconvolution [15] to cluster these metagenomic contigs into putative MAGs (supplementary information). This yielded >1000 clusters of contigs for each sample, of which 51 (WW) and 38 (WWEC) were >80% complete bacterial MAGs, as measured by CheckM [17] (Fig. S2, Tables S1-S2). In this paper, we use the term “cluster” to describe a cohesive group of contigs belonging to a genome of a microorganism. The EC genome was represented by one large cluster (4.2 Mbp) and three small ones (480 Kbp total) with similar high abundance, producing a >97% complete *E. coli* genome (Fig. S3A). Furthermore, Hi-C linkage between pB10::rfp and its host was extremely strong relative to other clusters (Fig. S3B), confirming that Hi-C can accurately ascertain plasmid-host relationships within a natural diverse microbial community.

### Cultivation-independent identification of host-ARG and host-plasmid associations

We investigated if Hi-C links could be used to identify the hosts that contain ARGs, integrons, and plasmids directly from the WW sample. We searched the WW and WWEC metagenome assemblies for sequences demonstrating high similarity to described plasmids [18], ARGs [19], or integrase genes (Tables S3-S4). Additionally, we inferred phylogenomic placements of each cluster. We then computed the Hi-C linkage of each ARG-bearing or plasmid-bearing contig to each cluster (Fig. 1 and Figs. S4-S8). ARGs were mostly linked to contigs in clusters related to the Gammaproteobacteria, Betaproteobacteria and the Bacteroidetes (Figs. S5-S6). Reproducible linkages observed in both WW and WWEC samples pointed to specific candidates in this wastewater sample that are well known to carry these ARGs. For example, clusters related to *Prevotella* and *Bacteroides* were linked to *tetQ, ermG, mefA, bla*_CFX_ and *bla*_CBLA_ conferring resistance to tetracycline, macrolides/lincosamides/streptogramins (MLS), and beta-lactams, respectively (Fig. S7)[20–23]. Similarly, in the *Acinetobacter*, the ARGs *mphE* and *tet39* were widespread in several clusters affiliated to the genus [24], and an *ant3*’’ and class 3 integron were linked with a cluster related to *Acinetobacter johnsonnii* [25]. Strikingly, the bacterial taxa identified to have the most contacts with known ARGs were affiliated with the *Aeromonadaceae* (Fig. 1-A), a taxonomic group typically associated with aquatic environments. Overall, the results are in line with current knowledge of the taxonomic distribution of these ARGs. This technique suggests that *Aeromonadaceae, Acinetobacter, and Bacteroidetes* are reservoirs of ARGs in this wastewater, as also recently suggested for another wastewater sample in Portugal using different approaches [26].

**Fig. 1:**
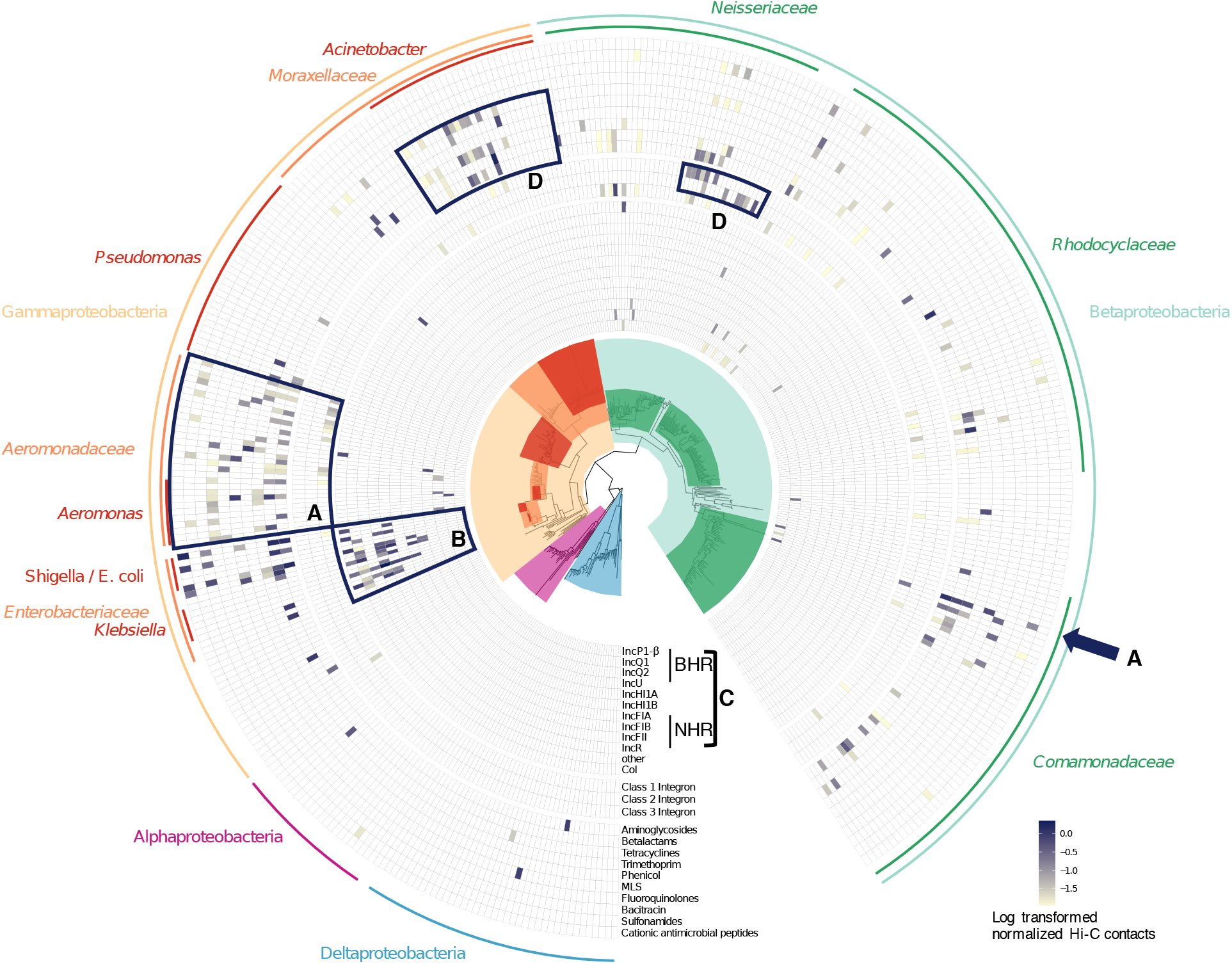
Hi-C linkage between plasmid markers, integrons, and ARGs among clusters belonging to Alpha-, Beta-, Gamma- and Deltaproteobacteria in the wastewater sample WW. Clusters are arranged in the inner circular phylogenetic tree where each tip represents a cluster. The presence or absence of a link is shown in the heatmap circling the tree, with the shading representing the intensity of the normalized Hi-C linkage signal. **A)** Identified natural reservoirs of ARGs: *Aeromonadaceae* linked to 18 ARGs conferring resistances to eight antibiotic classes, as well as the class 1 integron integrase gene and plasmids of the incompatibility groups IncQ and IncU (originally described in *Aeromonas* [30]), and a col plasmid [31]; a cluster in *Comamonadaceae* linked to an IncP-1β plasmid, class 1 integron, and 11 ARGs (links suggest some ARGs are plasmid-borne, SI). **B)** Most plasmids detected belonged to clusters related to *Enterobacteriaceae*. **C)** As expected, markers for broad-host-range (BHR) plasmids were linked to clusters spanning both *Beta-* and *Gammaproteobacteria*, specifically the *Enterobacteriaceae, Aeromonadaceae, Neisseriaceae, Rhodocyclaceae*, and *Comamomadaceae*. In contrast, markers for narrow-host-range (NHR) plasmids were almost exclusively linked to clusters belonging to the *Enterobacteriaceae*. Only one contig with a marker for an IncFIB plasmid was found in a *Betaprotebacteria* cluster, but the link was ~100 times weaker than linkage with the *Enterobacteriaceae*. **D)** Clusters affiliated with the genus *Acinetobacter* showed high Hi-C linkage to the ARGs conferring resistance to aminoglycoside (*aacA3*), betalactam (*bla*_OXA_ genes), tetracycline (*tet39*), phenicol *floR*) and macrolides (*mphE*). Class 2 and 3 integron integrase genes were associated with clusters affiliated with the *Neisseriaceae* while the class 1 integrons had links to 39 clusters within the Beta- and Gammaproteobacteria. Class 2 integrons were previously found in *Neisseria* sp. isolated from a WWTP, but this may be the first time that class 3 integrons are found in this family (see caption of Fig S4).

We also compared the *in situ* host range of both broad-host-range (BHR) and narrow-host-range (NHR) plasmids (Fig. 1) [27]. The results were strikingly consistent with our expectations, with IncQ plasmids showing the broadest range of putative hosts, followed by the IncP-1β plasmids, which were widespread but limited to the Betaproteobacteria [27]. It should be noted that most plasmids were linked to clusters related to the *Enterobacteriaceae* (Fig. 1-B), which may be due to the bias in the plasmid database we used [18]. Nevertheless, we show here that the Hi-C method allows assessing the *in situ* plasmid host range without any cultivation steps (Fig. 1-C).

Interestingly, in the WW sample, one *Comamonadaceae* cluster with high genome completeness (88.4%; cluster.20) showed strong linkages to an IncP-1β plasmid, a well-known host-plasmid association [28, 29]. We were able to reconstruct two large fragments (22.7kb and 12.9kb) carrying the typically conserved transfer regions of IncP1-ß plasmids and genes of the maintenance/control region. The closest relative was plasmid pALIDE02 of a WWTP isolate, *Alicycliphilus denitrificans* BC of the *Comamonadaceae* (see SI for some limitations of the method). Among the integrons, class 1 integrons exhibited links to the most clusters, and were the most widespread among the Proteobacteria. We conclude that the Hi-C links can help in determining the taxonomic placement of the hosts of ARGs and mobile elements in an environmental habitat despite some methodological challenges to be addressed in future work (see SI).

### Clustering-independent identification of plasmid and ARG hosts

It is possible that ProxiMeta has misclustered contigs, giving us chimeric clusters composed of contigs from different microorganisms. To independently verify the accuracy of our plasmid/integrons/ARGs host assignments, we performed taxonomic profiling of all the contigs that were linked to plasmid, integrons, and ARGs contigs. Fig. 2 shows that the host taxonomy of the contigs linked to these genes and mobile elements were mostly similar to those identified by ProxiMeta (Fig.1).

**Fig. 2:**
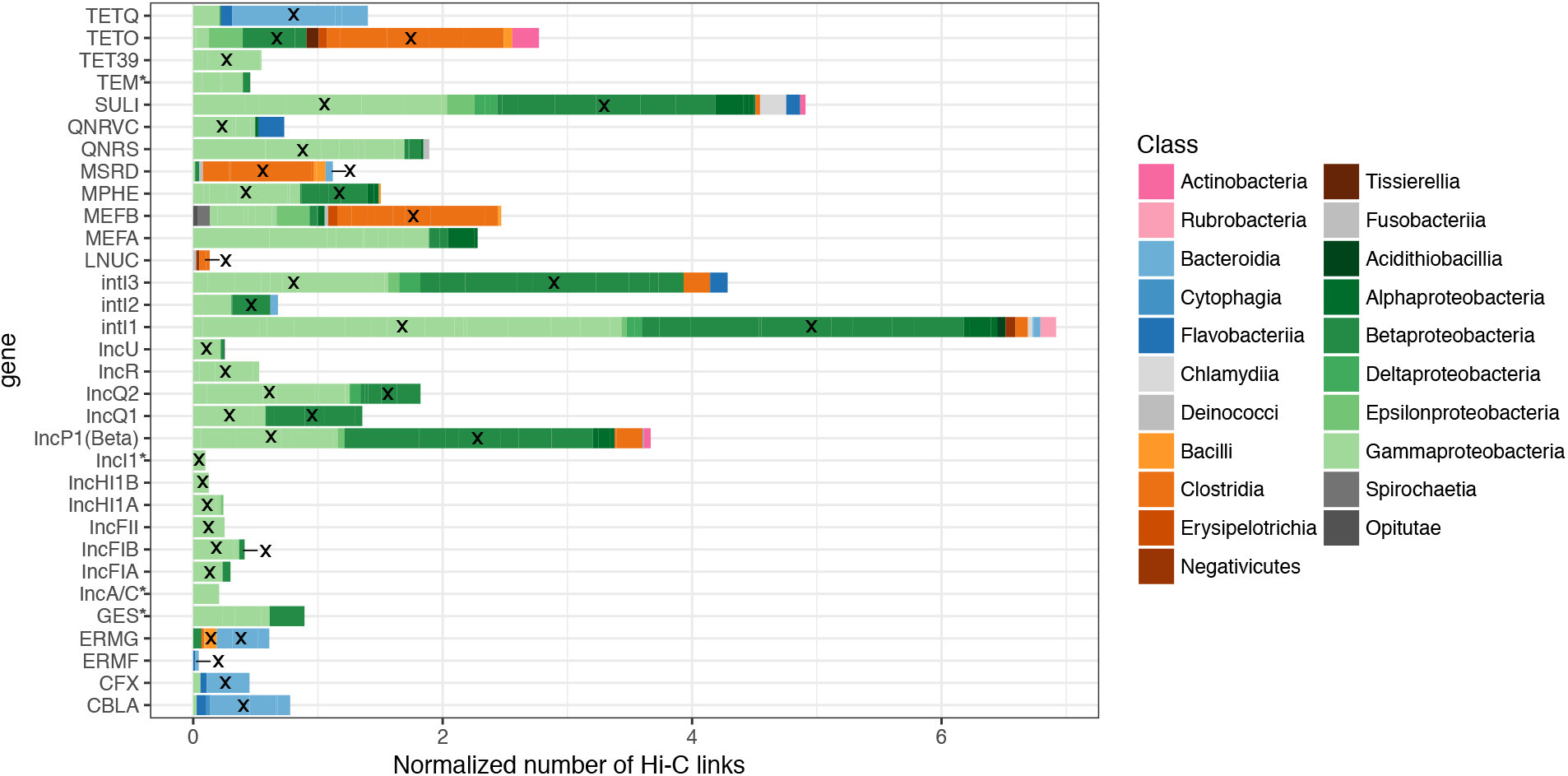
Taxonomic assignment of contigs that were linked to contigs harboring plasmids, ARGs, or integrons. We selected contigs harboring either a plasmids marker, an ARG, or and integrons integrase gene, and isolated all the contigs linked to these contigs by at least one Hi-C links. We used BLAST to find matches of each such linked contig against the NCBI bacterial database (see Materials and Methods, SI). Here the strength of Hi-C linkage is represented by the length of the bars and is summarized by Phylum (pink, Actinobacteria; blue, Bacteroidetes; orange, Firmicutes; green, Proteobacteria, and grey: others), by Class (color shading), and stacked bars of the same color represent different Families. *: ARGs or plasmids which did not have links to cluster when using our first approach. The cross indicates that the gene, or marker, was found in the same class when using our first approach (Fig. 1). For example, the MLS resistance gene *mphE* was strongly associated with contigs related to both Gamma- and Betaproteobacteria. More specifically links were associated with contigs belonging the *Moraxellaceae* and *Neisseriaceae*. Also, both quinolone resistance genes *qnrS* and *qnrVC* were mainly associated with contigs of the Gammaproteobacteria, mostly related to the *Aeromonadaceae*. As in our previous approach, markers for BHR plasmids were linked to clusters spanning both Alpha-, Beta- and Gammaproteobacteria while markers for NHR plasmids were strongly linked to clusters belonging to the Gammaproteobacteria, in particular the *Enterobacteriaceae*. Only the phylogenetic attribution of the ARG *mefA* was different between our two approaches.

## Conclusions

While several questions about the accuracy and sensitivity of this approach remain, we show that *in vivo* proximity ligation can help assess the *in situ* host range of ARGs, plasmids, and integrons in a natural microbial community. Such analysis can be expanded to other mobile genetic elements and other habitats with complex communities. This novel approach fills an important gap in our ability to track the reservoirs and horizontal transfer of antibiotic resistance genes, with the ultimate goal of slowing down the spread of drug resistance.

## Supporting information

## ACKNOWLEDGEMENTS

Research reported in this publication was supported by an Institutional Development Award (IDeA) from the National Institute of General Medical Sciences of the National Institutes of Health under grant number P30 GM103324 through an IBEST Pilot Grant, and in part by grants 2R44AI122654-02A1 and 1R43AI122654-01 to Phase Genomics. We thank the IBEST Genomics Resources Core staff Dan New, Matt Fagnan and Sam Hunter for assistance with shotgun library preparations and sequencing.

## CONTRIBUTIONS

TS, ET, and IL conceived the project, TS and MP wrote the manuscript, IL and ET provided thorough revisions, TS collected the sample and prepared the libraries, MP, SS, IL and TS did the bioinformatics analysis.

