## Supplementary material for "Linking the Resistome and Plasmidome to the Microbiome"

#### Materials and Methods

##### Sample collection

The wastewater sample was collected in October 2017 at the Moscow wastewater treatment plant (WWTP) in Idaho (USA). The facility services approximatively 25,000 people and collects mainly domestic wastewater. The sample was collected at the entrance of the WWTP, and 2 ml subsamples were centrifuged at 12,000 g for 3 minutes. Pellets were archived at -20°C until further processing.

##### Sample processing

Wastewater pellets were thawed on ice. In half of the aliquots we added a known number of *E. coli* K12::gfp containing plasmid pB10::rfp cells [1], hereafter named EC, such that they represented approximately 10% of the total bacterial community. Wastewater aliquots spiked with EC are named WWEC and aliquots consisting of the wastewater only are named WW.

The EC strain was grown overnight at 37°C from a glycerol stock (-70°C) in LB broth containing nalidixic acid (50mg.L$^{-1}$), kanamycin (50mg.L$^{-1}$), and tetracycline (10mg.L$^{-1}$). The culture was centrifuged for 3 minutes at 5,000g and pellets were resuspended with the same volume of PBS. We estimated the number of bacteria in the wastewater and in the *E. coli* culture using three methods as described below.

First a qPCR approach was used to target the 16S rRNA gene [2]. Reactions were carried out in a final volume of 10µL, with 1µL of pure and 10-fold diluted total genomic DNA from the WW or the bacterial culture, and the PerfeCta® qPCR ToughMix 2X (Quanta BioSciences™). The assays were performed in duplicate with an Applied Biosystems StepOnePlus Real-Time PCR System (Applied Biosystems™), and a standard plasmid containing the targeted sequence was used to construct a full standard curve in duplicate. The total bacterial cell count in the *E. coli* culture was estimated by dividing the number of 16S rRNA gene copies by the gene copy number in *E. coli* K12 (7). The total bacterial cell count in the water sample was estimated by dividing the number of 16S rRNA gene copies by 4.9, the average 16S rRNA gene copy number among 10,996 bacteria in the rrnDB database (https://rrndb.umms.med.umich.edu/). The WW sample contained an estimated number of 3.5 x $10^8$ bacteria/$ml^{-1}$ and the *E. coli* culture 2.7 x $10^9$. Using flow cytometry and viable plate counts on R2A agar, the estimated bacterial counts were very similar: 3.3 x $10^8$ and 2.7 x $10^7$ per ml respectively for WW and 3.5 x $10^9$ for the *E. coli* culture with both methods.

For the pure culture, total genomic DNA was extracted from 2 ml of the overnight culture using the GenEluteTM Bacterial Genomic DNA kit (Sigma-Aldrich, St. Louis, MO, USA).

**Library preparation**

Shotgun metagenomic libraries were prepared as follows. Total genomic DNA was extracted and isolated from WW and WWEC pellets using the DNeasy® PowerWater® kit (Qiagen). PCR-free Illumina libraries for short insert length sequencing with Hiseq were made by the IBEST Genomics Resources Core (Moscow, ID, USA) using TruSeq® DNA PCR-Free library Prep kit (Illumina). The Hi-C libraries were prepared from the WW and WWEC pellets using the ProxiMeta™ Hi-C preparation kit (Phase Genomics).

**Sequencing**

We pooled the Hi-C and shotgun metagenomic libraries, and sequenced both samples using two lanes of HiSeq 4000, 2x150bp paired end reads at the University of Oregon sequencing core

(Eugene, OR, USA). This produced 269,312,499 and 95,284,717 read pairs for the WW shotgun metagenomic and Hi-C libraries, respectively (ratio Hi-C:shotgun = 0.35), and 291,051,995 and 117,388,834 read pairs for the WWEC shotgun metagenomic and Hi-C libraries (ratio Hi-C:shotgun = 0.40).

**Data processing**

Hi-C data analysis was performed using the ProxiMeta workflow as previously described [3]. Some steps are described in more detail below.

*Processing of the shotgun sequencing data.* Shotgun sequencing data were processed using the following steps. Sequencing adapters were removed using BBDuk (BBTools developed by the Joint Genome Institute) with options k=23, ktrim=r, mink=12, hdist=1, minlength=50, –tpe, –tbo. Low-quality bases will be trimmed with BBDuk and options qtrim=rl, trimq=10, minlength=50, chastityfilter=True.

*Metagenomic assemblies.* Shotgun metagenomic assemblies were created *de novo* using Megahit with default parameters [4]. *De novo* assemblies were assessed using MetaQuast [5].

*Processing of the Hi-C reads.* Each set of reads was mapped to the metagenomic assemblies. Mapping was done using the Burrows-Wheeler alignment tool bwa mem [6, 7]. All reads that were incorrectly paired, unmapped, not uniquely mapped, mapped with a MAPQ score less than 20, or read pairs mapping to the same contig (which are not informative for deconvolution) were removed from the analysis.

*Deconvolution of the Hi-C data:* Deconvolution of contigs in the *de novo* assembly was performed using the ProxiMeta platform [3] that is partly based on the previously described [8].

Briefly, contigs less than 1000 bp in size, or which contained fewer than two restriction sites for the relevant enzyme were discarded for purposes of clustering. This dataset was normalized by the number of restriction sites on the contigs and contig Hi-C read coverage (which implicitly accounts for length and abundance, among other characteristics). Finally, the contigs were grouped into clusters based on their Hi-C linkages using a proprietary Markov Chain Monte Carlo algorithm.

*Annotation of genome clusters.* Genome clusters were compared to RefSeq genomes using Mash [9] to identify any close database matches for new genome clusters. Genome clusters were further analyzed using the CheckM [10] lineage_wf workflow with the --reduced_tree option to assess genome quality and estimate high-level phylogenetic placements for each cluster based on single-copy marker gene analysis. In some cases, additional manual merging of genome clusters was performed based on sequence characteristics and CheckM completeness/contamination criteria. Some genome clusters were excluded on the basis of promiscuous interactions with other clusters, quantified as their vertex entropy in the Hi-C graph connecting genome clusters, calculated using the R entropy package [11]. We excluded clusters with vertex entropy higher than 3. We assessed abundance of different organisms as the median of the abundance of constituent contigs greater than 20Kb in size, estimated as kallisto "transcripts per million" [12]. Circos plots were generated using Circoletto [13], and genome alignments to references were analyzed using quast v5.0.0 [14].

*Marker gene detection in the metagenome assemblies.* Detection of marker genes was done using BLAT with the options -minIdentity=90, and hits for which the coverage of the reference sequence was lower than 80% were discarded. When a contig had multiple hits for the same locus, we selected the best hit (best score obtained by multiplying the coverage by the identity). Detection of the antibiotic resistance genes and plasmids were respectively done using the MEGARes database [15] and the PlasmidFinder database [16], accessed in April 2018. The MEGARES database was depleted from all genes for which resistance is conferred by SNPs, multi-drug efflux pumps, and regulators. Detection of the class 1, 2 and 3 integron integrase genes was done using the reference sequences AB709942 (*intI1*), FQ482074 (*intI1*delta1), JX566770 (*intI1*R32_N39 aa329-337 mutated + 35aa), JX469830 (*intI2*), EF467661(*intI3*) [17].

*Linking plasmid, ARG, and integron contigs to genome clusters.* We used a simple heuristic to infer linkages between genome clusters and contigs thought to carry ARGs or plasmid sequences. For each sample, we considered all Hi-C linkages between contigs of interest and any other contig. We then discarded all contig-cluster linkages represented by only one or two reads. We next used the same criteria to infer all linkages between ARG and plasmid contigs. Phylogenetic analysis of plasmid-genome interactions was performed using the CheckM tree workflow and ape v5.1 [18] and phytools v.0.6-44 [19]. We normalized the number of Hi-C contacts (reads) according to abundance of the plasmid/ARG and abundance of the genome cluster. We discarded contacts <0.01 as spurious. We then summarized all normalized contacts across all contigs corresponding to each plasmid, integron, or ARG family (multiple contigs can correspond to the same plasmid, integron, or ARG family).

*Data availability:* Sequencing data are available in FASTQ format at SRA accession PRJNA506462. Processed data and scripts for linking contigs to genome clusters using Hi-C data are available at https://osf.io/ezb8j/.

*Taxonomic summaries of Hi-C linkages with plasmid/ARG/integron contigs.* As an alternative measure of plasmid/ARG/integron - host interactions that did not depend on the accuracy of genome clusters, we ignored ProxiMeta clustering results and considered only the set of contigs to which each contig of interest (i.e. plasmids, ARGs, or integrons) is connected by Hi-C. We used BLAST to search these contigs, considering only the best hit among hits with high confidence (E-value < 1e-20). We mapped these hits to NCBI taxonomic identifier (taxids) and species names where possible, otherwise recording "no hit" as the species name. We then filtered out contigs for which the coverage of the alignment was <80%, contigs for which there was "no hit", and contigs identified as "uncultured" or "*Candidatus*" bacteria. Finally, we counted the number of Hi-C links to each species name, correcting for contig abundance. Where measured abundance was zero, we replaced it with a small nonzero number (0.1 transcript per million (TPMs)).

#### Intracellular association of ARGs in cluster.20

For the WW sample, we extracted the subgraph of Hi-C contacts between the 1497 contigs within cluster.20 (including plasmid, integron, and ARG contigs that had Hi-C contacts but were not clustered to cluster.20). We used Louvain clustering of this subgraph to identify contig components of the Hi-C graph that are associated with plasmids, integrons, and ARGs (figure below). We found that all but one of the 13 plasmid, integron, and ARG contigs in cluster.20 are co-located in one Louvain subcluster of 31 contigs. To further identify contiguous sequences within this subcluster we did a *de-novo* assembly of the set of contigs using Geneious 8.1.9 (assembler: geneious using preset parameters "high sensitivity"). Seven of the 31 contigs produced a 22.7kb and 12.9kb fragment carrying all typically conserved transfer regions of IncP1-β plasmids (*tra* and *trb* genes) and genes of the maintenance/control region (pink circles on figure below). Of the 10 ARGs, $bla_{\mathrm{OXA9}}$ and *ant*(3'') did not have a direct Hi-C link with a contig harboring a conserved region of IncP1-β plasmids suggesting they were not located on a IncP-1β plasmid. The rest, *aph(3')Ia*, *tetA*, *aph(3'')Ib-aph(6)-Id*, $bla_{\mathrm{OXA2}}$-*ant*(3'')Ia, *tetC*, *sul1* and *catA* had a direct Hi-C link with contigs harboring a conserved region of IncP-1β plasmids suggesting that they were plasmid borne.

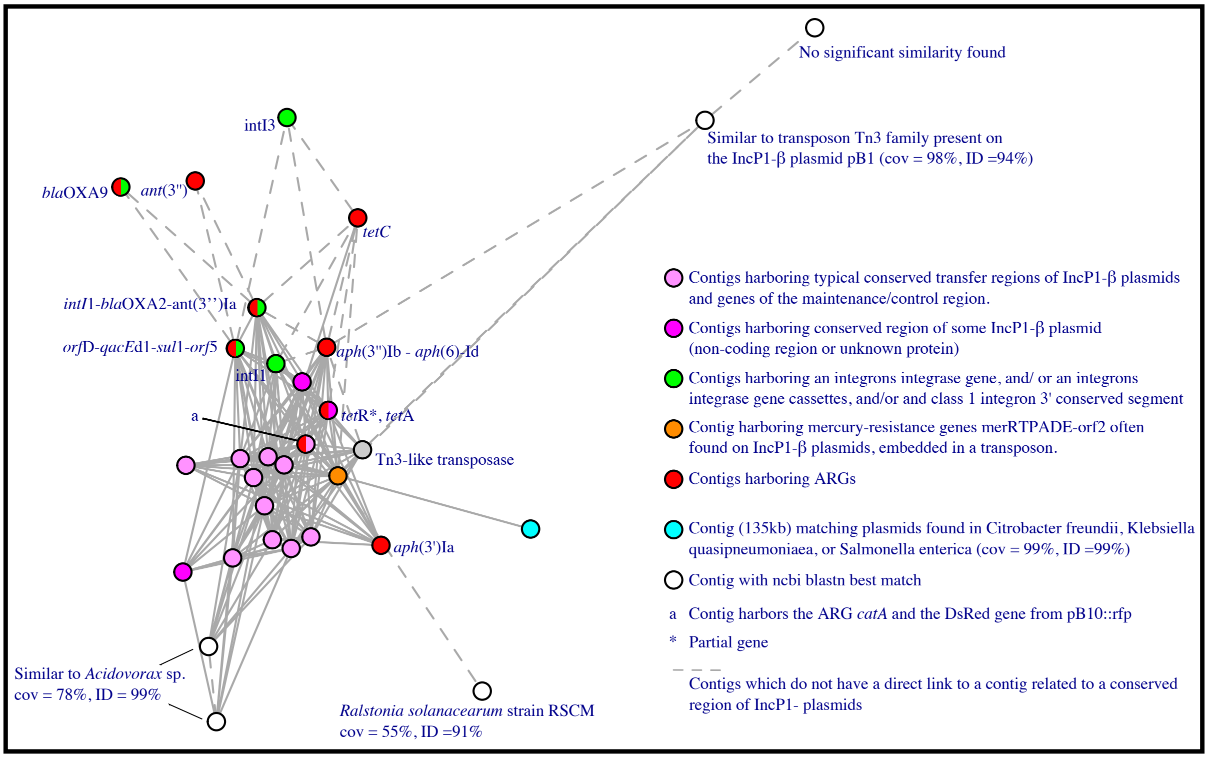


Surprisingly, one of the plasmid contigs contained the *dsRed* gene and the *catA* resistance gene used to mark the plasmid pB10, resulting in pB10::rfp. This plasmid was present in the EC strain used to spike the WWEC sample but should not be present in the WW sample. Consistent with that *dsRed* was not detected by qPCR in the total WW gDNA used to make the shotgun library (data not shown). These findings suggest some level of cross-contamination between the WW and WWEC libraries. The *gfp* gene present in the chromosome of the spiked EC strain was not linked to a cluster in our clustering analysis, and we detected very few Hi-C links between clusters related to *E. coli* and genes found on pB10::rfp. These results show that the cross-contamination was minor. Since here the samples WW and WWEC were the same, such cross-contamination does not affect the overall interpretation of our result. However, it affects our ability to determine whether the ARGs linked to cluster.20 were present in the *Comamonadaceae* or belonged to pB10::rfp. Indeed, seven of the nine potential plasmid-borne ARGs in cluster.20

were the ARGs carried by pB10::rfp. Nevertheless, the presence of IncP-1beta plasmids in this cluster is real, as the 22.7kb fragment carrying the *tra* genes of IncP1-β plasmids was identical (>99% coverage >99% identity) to 18 IncP1-β plasmids other than pB10, and the 12.9kb fragment carrying the *trb* genes of IncP-1β plasmids was identical (>99% coverage and >99% identity) to 33 different IncP1-β plasmids, including pB10. The closest relative to both fragments was the plasmid pALIDE02 of *Alicycliphilus denitrificans* BC, a *Comamonadaceae* isolated from WWTP [20]. Members of the *Comamonadaceae* and more broadly the Burkholderiales are well-known hosts of IncP1-β plasmids[21–23]. We concluded that the WW sample contained native IncP1-β plasmids present in *Comamonadaceae* but that many of the ARGs detected may come from the pB10::rfp cross-contamination. This result highlights one of the limitations of the approach when it comes to resolving the location of mobile genetic elements shared among different bacteria (see section below, "limitations").

#### Limitations

*Highly abundant genomes may produce clustering artifacts*

In WWEC, several clusters showed Hi-C linkages to all or several genes naturally present in the chromosome of *E. coli* K12 (Fig. S6). As expected, those genes ($bla_{EC}$, *pmrA, B ,C, F, acrD*, *E*, *F*, *S*, *pbp4b*, *pbp2, ampH*, and *arnA*) were all on contigs linked to the EC cluster (black arrow in Fig. S6). However, the other clusters were spread out all over the phylogenetic tree and were related to the Firmicutes, Alpha- or Betaproteobacteria (open arrows in Fig. S6). The presence of these genes in such taxa is very unlikely. Besides, some of those genes were not detected in the WW sample suggesting that they were specific to the spiked EC. We conclude that the Hi-C linkages detected in the WWEC were spurious links probably related to the high abundance of the spiked organism.

*Genetic elements shared by several bacteria may produce spurious links*.

In WWEC, we observed Hi-C links between the EC cluster and ARGs or plasmid markers that do not belong to the genome of the spiked EC. For example, we detected Hi-C links to the markers of IncFIB, IncFII, and col plasmids, and to the ARGs *catB*, $bla_{OXA4}$, $bla_{OXA9}$, $bla_{TEM}$, $bla_{CTX}$. The presence of such plasmids or ARGs in other strains of *E. coli*, or closely related species, naturally present in the wastewater is very likely. This was supported by the fact that in WW several clusters related to *Escherichia* sp. and different from the strain EC were detected (Fig. 1 and Fig. S5), some of which had Hi-C links to these genes. We conclude that the presence of several related species or strains in the sample can produce *in silico* crosslinks between clusters that could not be resolved.

The same observation was made in our analysis of the cluster.20 (see above). Our result suggests that the cluster.20 contained a IncP1-β plasmid but that most of the ARGs having Hi-C links to this cluster may have come from the plasmid pB10::rfp, also a IncP1-β plasmids, due to minor cross-contamination between the WW and WWEC libraries. A similar observation was made for the cluster.1046, which shared some contigs closely related to *E. coli* and ended up to have Hi-C links to the ARGs from the spiked EC (see details in caption of Fig S8).

These limitations can be addressed by abundance normalization schemes such as those that we employ in this study, though these of course require accurate abundance estimates. As our understanding of Hi-C in metagenomics communities and the resulting numerical linkage data improve, we anticipate that these limitations will be addressed in future studies.

#### SUPLEMENTARY FIGURES LEGENDS

**Figure S1**: Hi-C deconvolution workflow. (A) Formaldehyde induces covalent bonds between DNAs that are close in three-dimensional space, and therefore within the same cell. (B) Hi-C library preparation via restriction enzyme digestion, religation, and junction enrichment reads out proximity information as links. (C) Hi-C links can be used in a graph clustering context to deconvolute contigs into their original cellular groupings, including both chromosomes and plasmids. (reproduced from Press *et al.* 2017 [3], reuse authorized under CC-BY-NC-ND license).

**Figure S2**: Completeness and contamination of each ProxiMeta genome cluster in each sample, as estimated by CheckM from single-copy marker genes. Current thresholds used to classify the quality of draft genomes recovered from metagenomic assembly, commonly named MAG (Metagenome Assemble Genome), were defined by the authors of CheckM [10]. The completeness of MAG and their associated brackets in percentage were defined to be “near” (≥90%), “substantial” (≥70% to 90%), “moderate” (≥50% to 70%), and “partial” (<50%). The contamination of MAG and their associated brackets in percentage were defined to be “low” (≤5%), “medium” (5% to ≤10%), high (10% to ≤15%), and very high (>15%).

**Figure S3**: (A) Alignment of the EC cluster to the reference genome of the *Escherichia coli* K-12 MG1655 genome and the plasmid pB10::rfp. Threshold for blast was $E<10^{-100}$. Coloring represents the percent pairwise nucleotide identity with blue ≤ 50%, green ≤ 79.9%, orange ≤ 89%.9, red > 89.9 % identity. The assembled genome cluster EC was composed of cluster.11 (44 contigs, 4.18Mb) and cluster.1069, cluster.925, and cluster.1037 (32 contigs, 478kb). Aligned against the reference genome sequence produced an alignment covering 97.5% of the reference covered with >99.9% identity. (B) Hi-C linkages faithfully recapitulate a known host-genome relationship between the plasmid pB10::rfp and its *E. coli* K-12 host spiked into the WW sample.

**Figure S4**: Hi-C links between, plasmid markers, integrons and ARGs among clusters that were identified to belong to Alpha- Beta-, Gamma-, and Delta-Proteobacteria in the WWEC sample. Clusters are arranged on the circular phylogenetic tree where each tip represent a cluster. The presence or absence of a link is shown on the heatmap circling around the tree, and the color shading represents the intensity of the normalized Hi-C link signal.

**A)** As in the WW sample, the *Aeromonadaceae* were shown to be a natural reservoir of ARGs, as clusters from this family linked to 22 ARGs, 14 of which were identical to the ones detected in the WW sample. Similarly, plasmids of the incompatibility group IncQ, and IncU were also associated with *Aeromonadaceae*. In addition, a marker for the IncA/C plasmid was detected in this family too. As for the IncU plasmids, IncA/C plasmids were first described in a *Aeromonas* species [24]. IncA/C plasmid markers were also detected in the WW sample but no links to clusters were detected. Here in WWEC no integrons were detected, possibly because the strong

Hi-C signal of the class 1 integron integrase gene from pB10::rfp. Finally, a marker of the IncP-6 plasmids not detected in the WW was linked to this family.

**B)** Fewer plasmid markers were detected in the WWEC compared to WW, but the markers detected also belonged to the IncFIB, IncFII, IncR, col and the other group of plasmids (i.e., Rep_1_pKPC-2_CP011573 and TrfA_1__CP11611).

**C)** Markers for BHR plasmids of the group IncQ were linked to cluster spanning both Beta- and Gamma-Proteobacteria and were related to the families of the *Enterobacteriaceae*, *Aeromonadaceae*, *Neisseriaceae*, *Rhodocyclaceae*, and *Comamomadaceae*. Here the marker for IncP1-β plasmids was only associated with the EC cluster, possibly because the links to pB10::rfp might have overwhelmed the analysis. On the other hand, markers for NHR plasmids were almost exclusively linked to cluster belonging to the *Enterobacteriaceae*.

**D)** As in the WW sample, clusters affiliated with the genus *Acinetobacter* showed high Hi-C linkage to the ARGs conferring resistance to aminoglycoside (*aacA3*), betalactam ($bla_{\mathrm{OXA}}$ genes), tetracycline (*tet39*), phenicol (*floR*) and macrolides (*mphE*). Similarly, the class 2 and 3 integron integrase genes were associated with clusters affiliated with the *Neisseriaceae*. More specifically, in both samples, the class 2 integron integrase was associated with clusters placed in the genus of *Neisseria* and the class 3 integron in the genus of *Vitreoscilla*. Class 2 integrons were previously found in *Neisseria* sp. isolated from a WWTP (Accession numbers FJ502342 and FJ502343). While a class 3 integrons were characterized from the *Delftia* sp. isolated from WWTP [25], they have never been found in *Neisseriaceae* (Integrall database, http://integrall.bio.ua.pt, consulted on 01/11/2018). This result suggest that this would be the first time that class 3 integrons are described in *Neisseriaceae*.

Finally, in the WWEC sample, Hi-C links between the class 3 integrase and the cluster.8 (completeness>90% and contamination <5%) were detected. The best taxonomic identification of this cluster was the family *Rhodocyclaceae*.

**Figure S5**: Hi-C links between all the clusters and plasmid markers, integrons, and ARGs, in the WW sample. The presence or absence of a link is shown on the heatmap to the right of the cluster tree, and the color shading represents the intensity of the normalized Hi-C link signals (Scale as in Fig. S3).

**Figure S6**: Hi-C links between all the clusters and plasmid markers, integrons, and ARGs in the WWEC sample. The presence or absence of a link is shown on the heatmap to the right of the cluster tree, and the color shading represents the intensity of the normalized Hi-C link signals (Scale as in Fig. S3). Arrows indicates clusters where Hi-C links suggest the presence of genes belonging to the spiked EC (see section "limitation"). The back arrow specifically shows the EC cluster.11.

**Figure S7**: Hi-C links between the clusters affiliated with the Bacteroides and ARGs in (A) the WW sample and (B) the WWEC sample. The presence or absence of a link is shown on the heatmap to the right of each tree and the color shading represents the intensity of the normalized Hi-C link signal. In both WW and WWEC clusters related to *Prevotella* and *Bacteroides* were linked to *tetQ*, *ermG*, *mefA*, *bla*$_{\mathrm{CFX}}$, and *bla*$_{\mathrm{CBLA}}$. The ARGs *tetQ,* and *ermG* or *ermF* are frequently found in tetracycline and erythromycin-resistant *Bacteroides* isolates, respectively [26, 27] (*ermF* was only detected in the WW sample). The *bla*$_{\mathrm{CFX}}$ has been found in both *Prevotella* and *Bacteroides* that produce beta-lactamases [28, 29]. *bla*$_{\mathrm{CBLA}}$ is species-specific and found in *Bacteroides uniformis* CARD database consulted Nov-29-2018 [30] . Finally, *mefA* and *bla*$_{\mathrm{CEPA}}$ (the latter one was only found in WW sample) are also commonly found in the *Bacteroides* [27].

**Figure S8**: Hi-C links within the Firmicutes between the clusters and ARGs, plasmid markers, and integrons in (A) the WW sample and (B) the WWEC sample. The presence or absence of a link is shown on the heatmap to the right of the tree and the color shading represents the intensity of the normalized Hi-C link signal. Several clusters showed a strong Hi-C signal to several contigs harboring a replication gene marker of a Gram-positive plasmid, "G+" in figures (reference sequence in the database was pKKS825). While in the WW those clusters were all affiliated with the family *Eubacteriaceae*, in the WWEC they were spread out over other families. The ARGs *ant9*, *tetO*, *cat*, *ermB*, *ermG*, *lnuC*, *mefA* and *mefB* were reproducibly found associated with clusters of the Firmicutes. Taxonomic affiliation of the cluster harboring *ant9* (*Lachtospiraceae*), *tetO* (*Lachtospiraceae*), *ermB* and *ermG* to (*Streptococacceae*), and *mefA* and *mefB* (*Proteocatella* sp.) was the same in both WW and WWEC.

Arrows indicates clusters where Hi-C links suggest the presence of genes belonging to the spiked EC (see section "limitation"). This cluster (cluster.1046) representing a total size of 46kb was comprised of 20 contigs for which 12 matched to the reference genome of *E. coli* K12 substr.

1655 with > 99% pairwise identity. The eight remaining contigs either mated the reference genome of *E. coli* K12 substr. 1655 with <99% pairwise identity, different strains of *E. coli* (for five contigs), or unknown or very distant bacteria. However here Proximeta did not cluster this small cluster with the rest of the *E. coli* K12 substr. MG1655 and this small cluster was therefore misplaced in the phylogenetic tree. Finally, one contig with a IncHI1A plasmid marker was linked to a *Firmicutes* cluster, an unlikely association as IncHI1A plasmids are thought to be found only in some Proteobacetria [31]; this was not observed in the WWEC sample.

#### SUPPLEMENTARY TABLES

**Table S1**: ProxiMeta report for WWEC sample

**Table S2**: ProxiMeta report for WW sample

**Table S3**: Contigs from WW sample identified to harbor an ARG, an integron integrase gene, or a plasmid marker.

**Table S4**: Contigs from WWEC sample identified to harbor an ARG, an integron integrase gene, or a plasmid marker.

**Table S5**: Quast reports for specific genomes.

#### SUPLEMENTARY FIGURES

### FIGURE S1

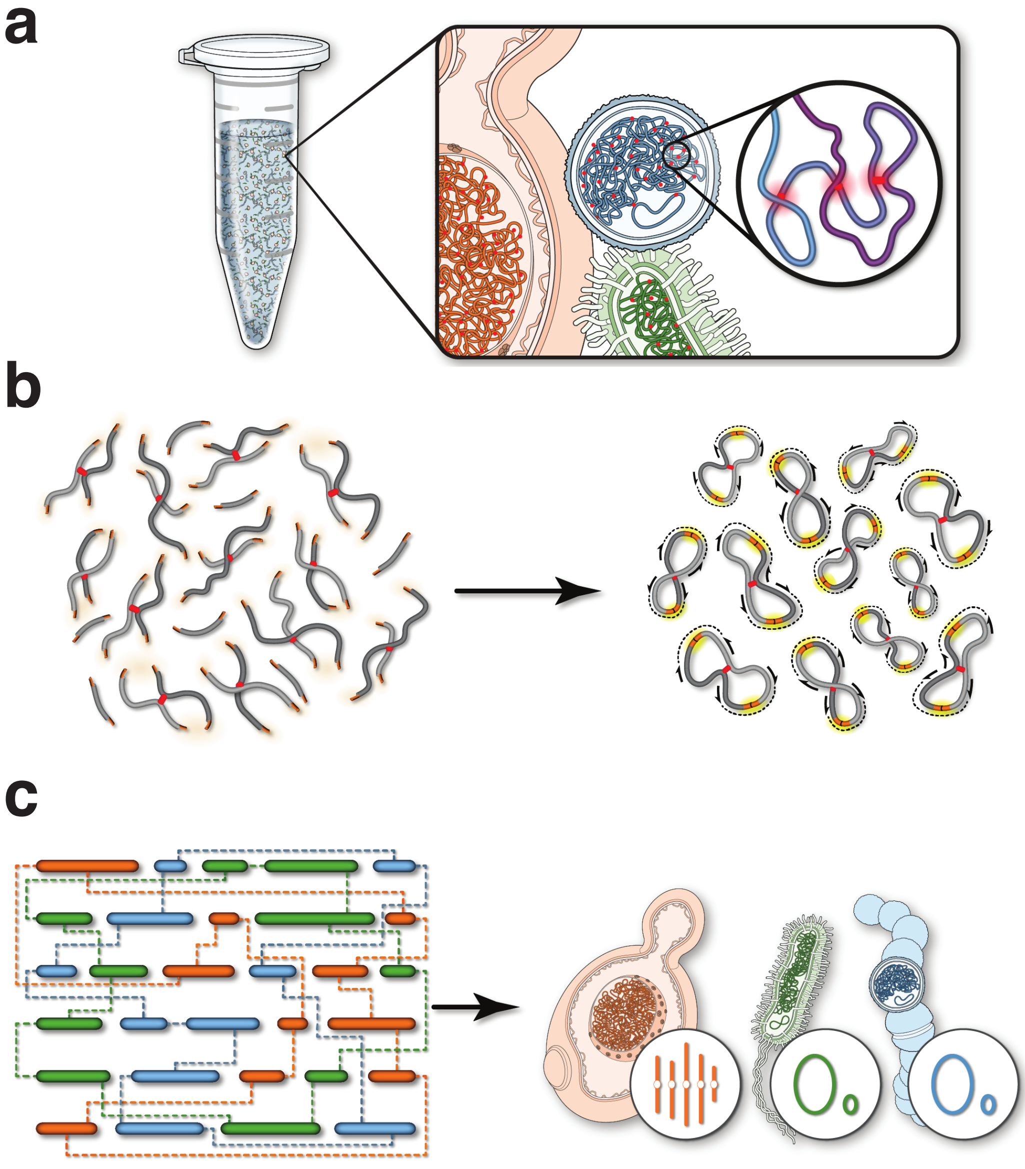

### FIGURE S2

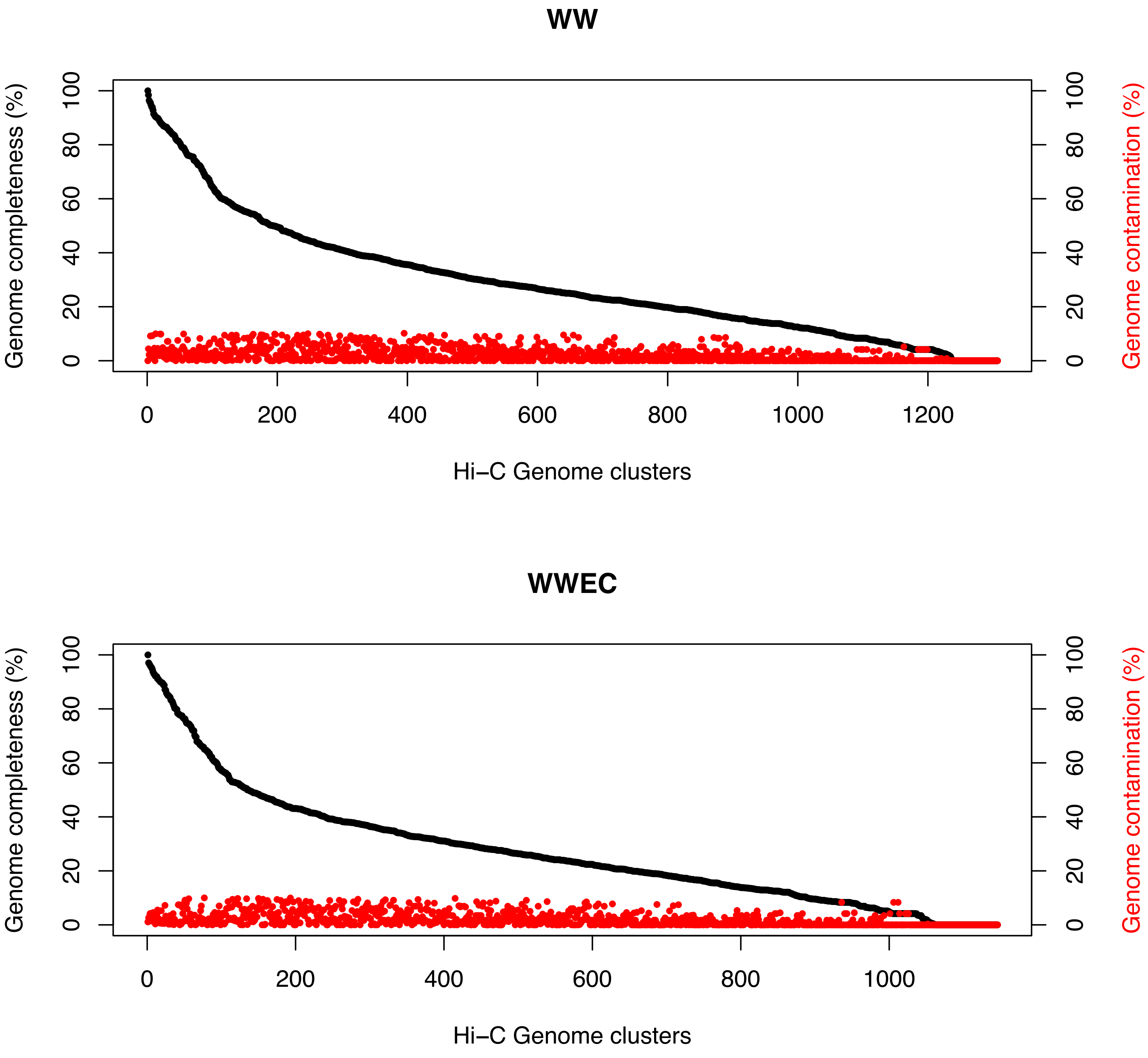
WW
Genome completeness (%)
Genome contamination (%)
Hi–C Genome clusters
WWEC
Genome completeness (%)
Genome contamination (%)
Hi–C Genome clusters

### FIGURE S3

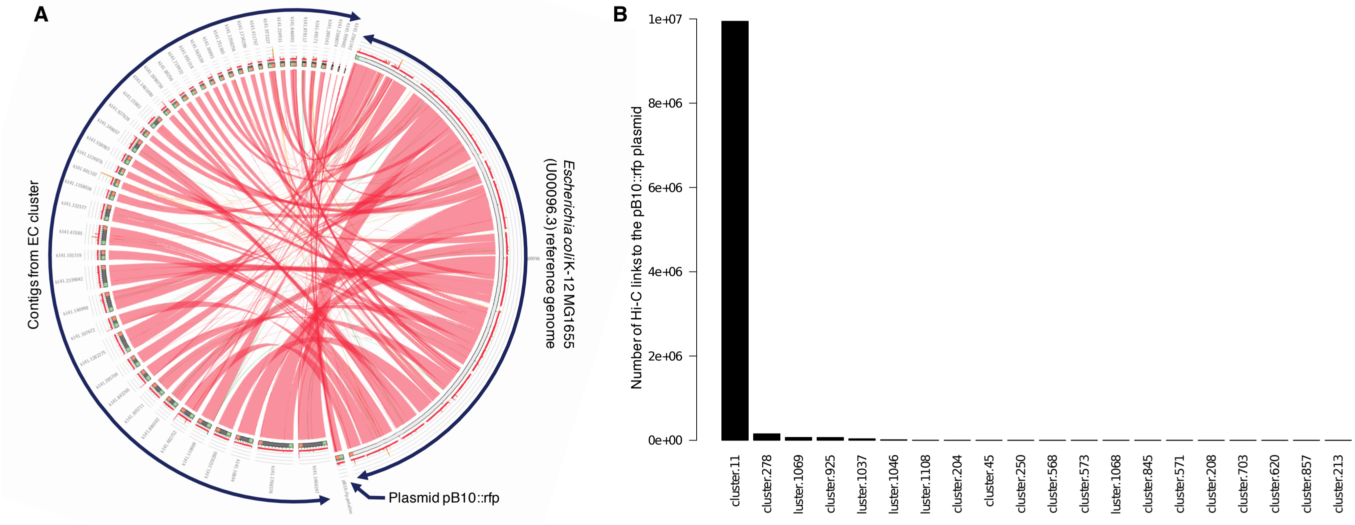
A
Contigs from EC cluster
Escherichia coli K-12 MG1655
(U00096.3) reference genome
Plasmid pB10::rfp
B
Number of Hi-C links to the pB10::rfp plasmid
1e+07
8e+06
6e+06
4e+06
2e+06
0e+00
cluster.11
cluster.278
luster.1069
cluster.925
luster.1037
luster.1046
luster.1108
cluster.204
cluster.45
cluster.250
cluster.568
cluster.573
luster.1068
cluster.845
cluster.571
cluster.208
cluster.703
cluster.620
cluster.857
cluster.213

### FIGURE S4

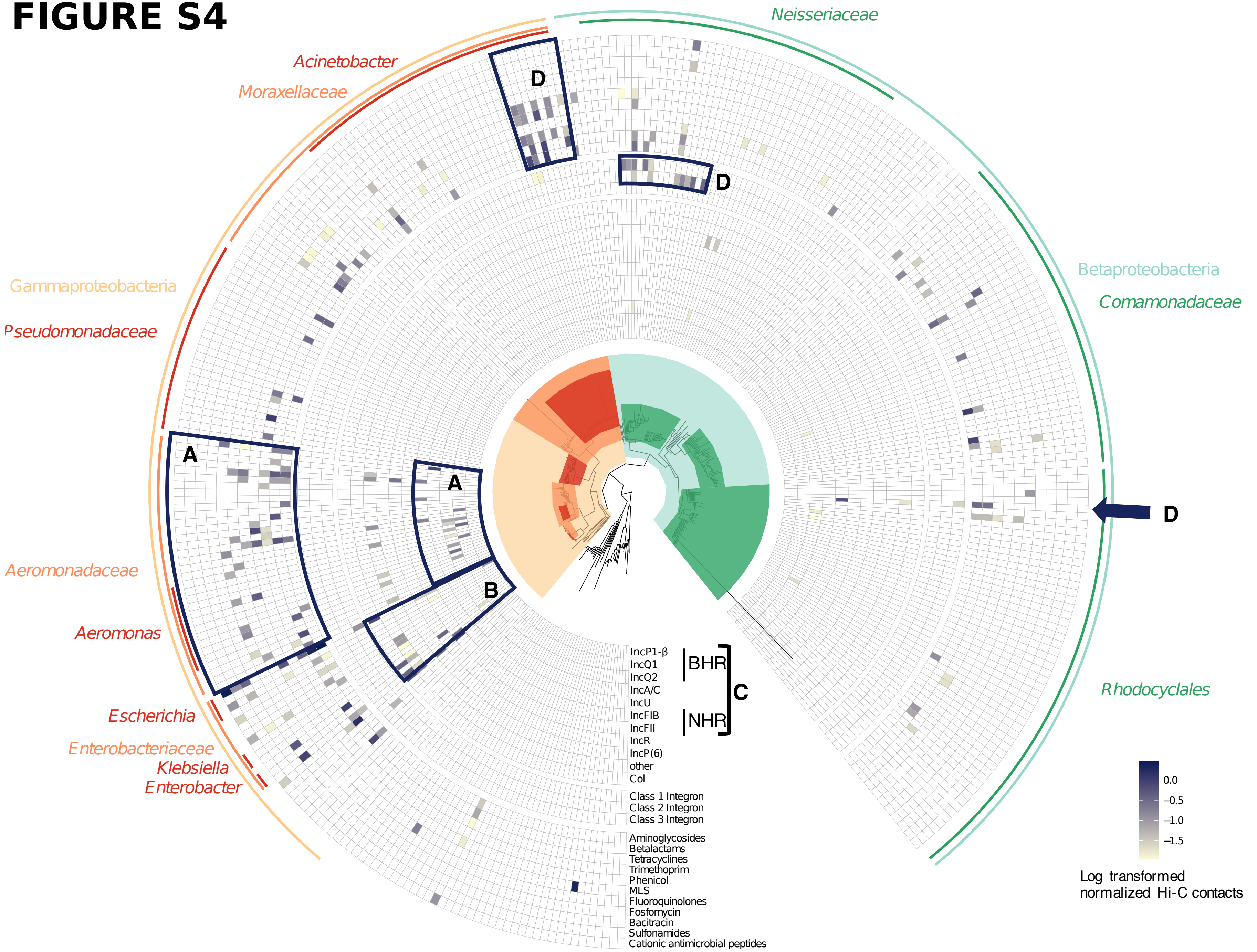
Neisseriaceae
Acinetobacter
Moraxellaceae
D
D
Gammaproteobacteria
Pseudomonadaceae
Betaproteobacteria
Comamonadaceae
A
A
D
Aeromonadaceae
B
Aeromonas
Escherichia
Enterobacteriaceae
Klebsiella
Enterobacter
Rhodocyclales
IncP1-β
IncQ1
IncQ2
IncA/C
IncU
IncFIB
IncFII
IncR
IncP(6)
other
Col
BHR
NHR
C
Class 1 Integron
Class 2 Integron
Class 3 Integron
Aminoglycosides
Betalactams
Tetracyclines
Trimethoprim
Phenicol
MLS
Fluoroquinolones
Fosfomycin
Bacitracin
Sulfonamides
Cationic antimicrobial peptides
0.0
−0.5
−1.0
−1.5
Log transformed normalized Hi-C contacts

### FIGURE S5

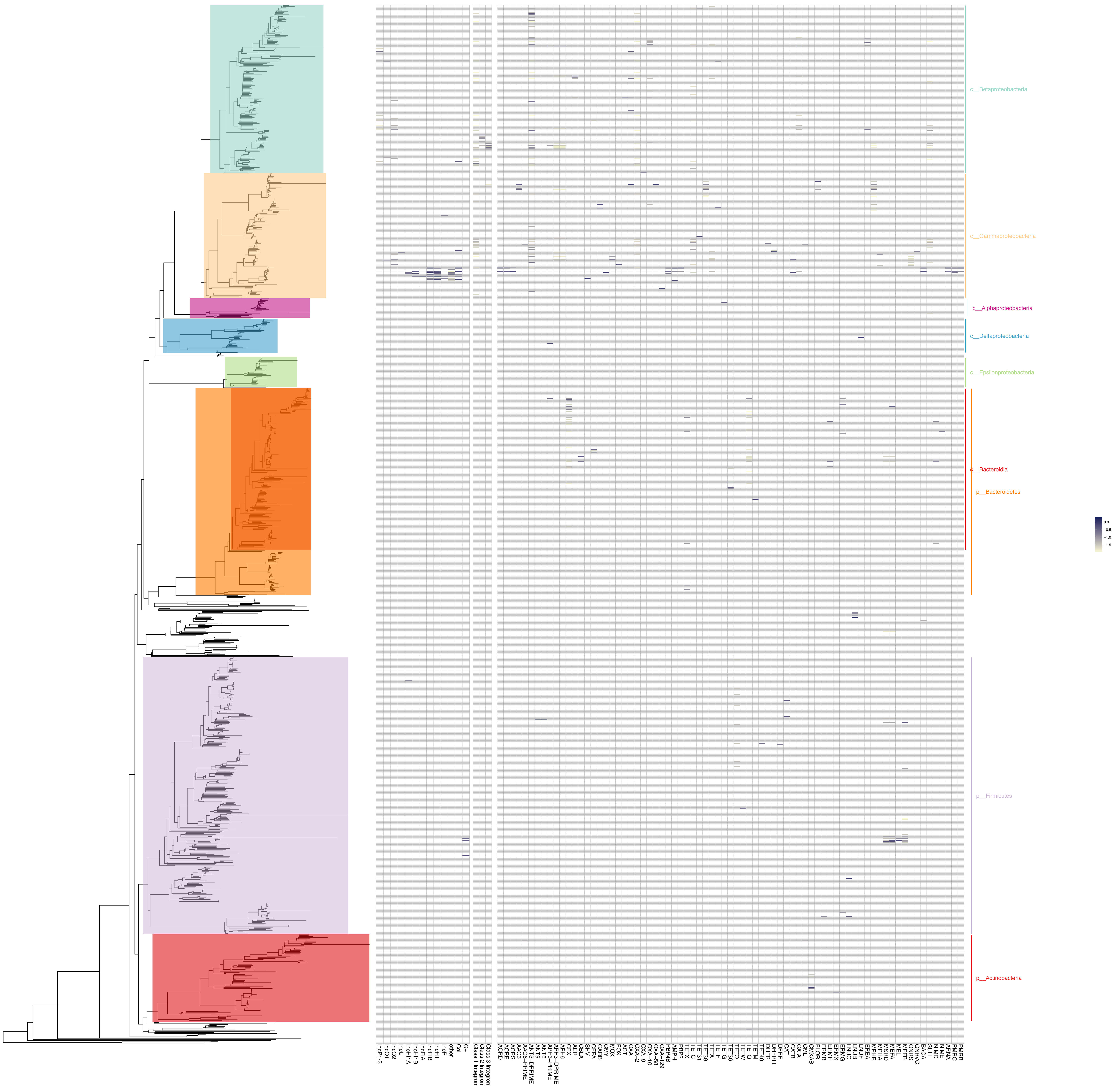
c__Betaproteobacteria
c__Gammaproteobacteria
c__Alphaproteobacteria
c__Deltaproteobacteria
c__Epsilonproteobacteria
c__Bacteroidia
p__Bacteroidetes
p__Firmicutes
p__Actinobacteria

### FIGURE S6

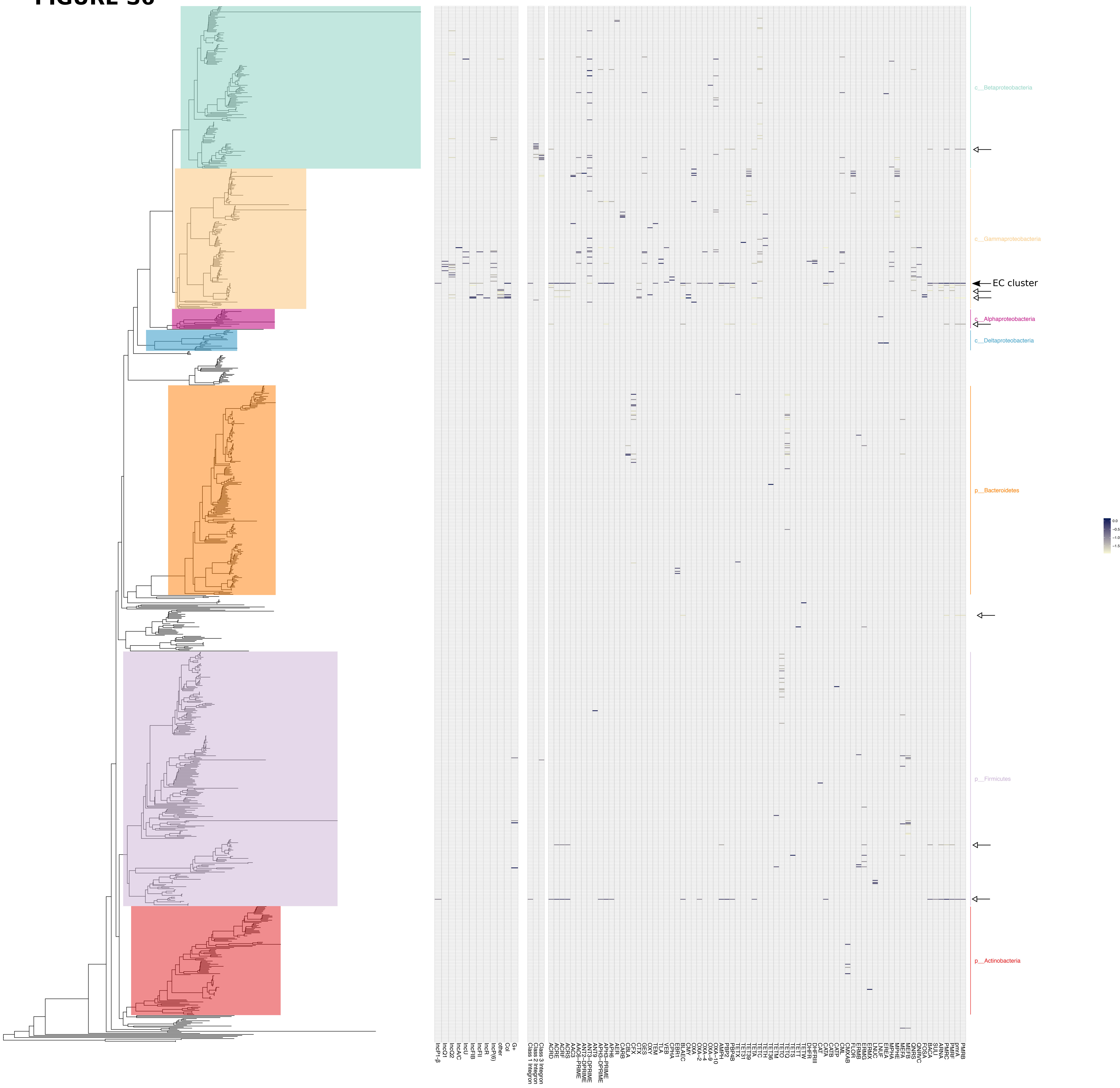

c__Betaproteobacteria
c__Gammaproteobacteria
EC cluster
c__Alphaproteobacteria
c__Deltaproteobacteria
p__Bacteroidetes
p__Firmicutes
p__Actinobacteria

### FIGURE S7

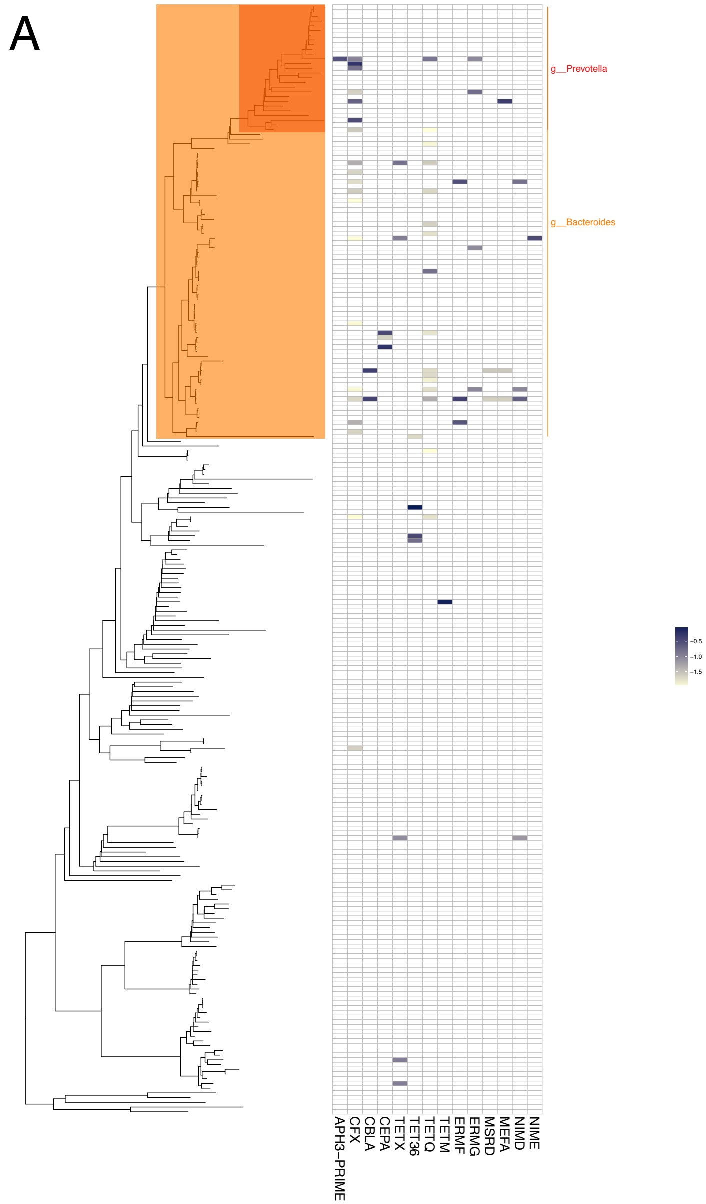
A
g__Prevotella
g__Bacteroides
APH3−PRIME
CFX
CBLA
CEPA
TETX
TET36
TETQ
TETM
ERMF
ERMG
MSRD
MEFA
NIMD
NIME
−0.5
−1.0
−1.5

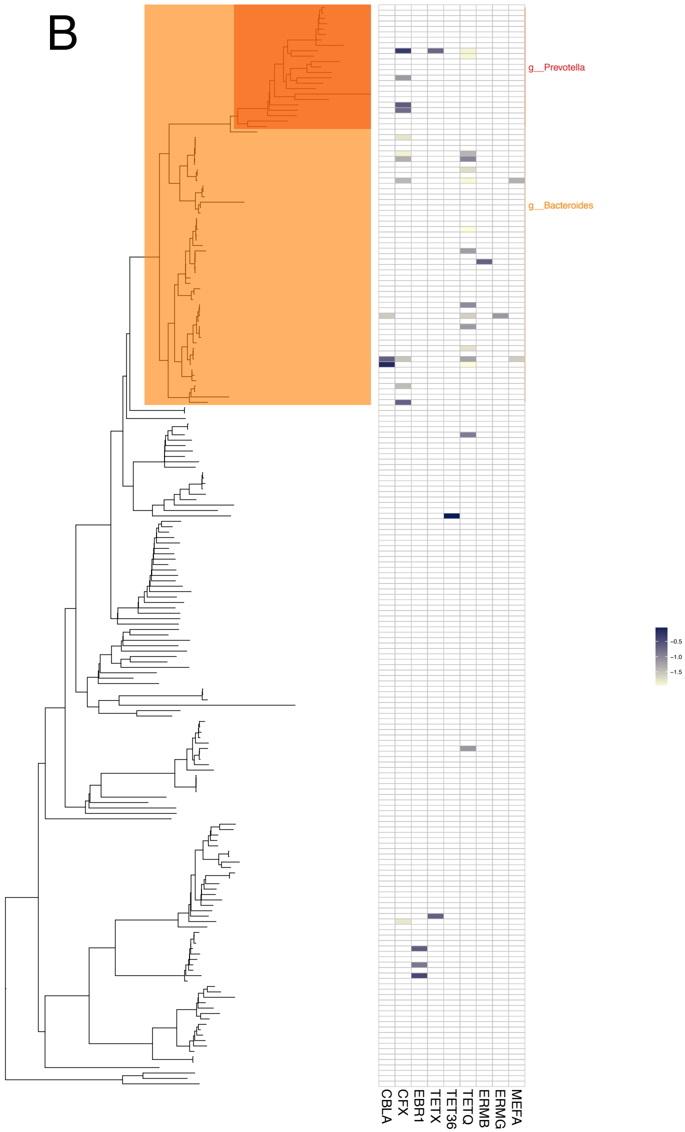
B
g__Prevotella
g__Bacteroides
CBLA
CFX
EBR1
TETX
TET36
TETQ
ERMB
ERMG
MEFA
−0.5
−1.0
−1.5

### FIGURE S8

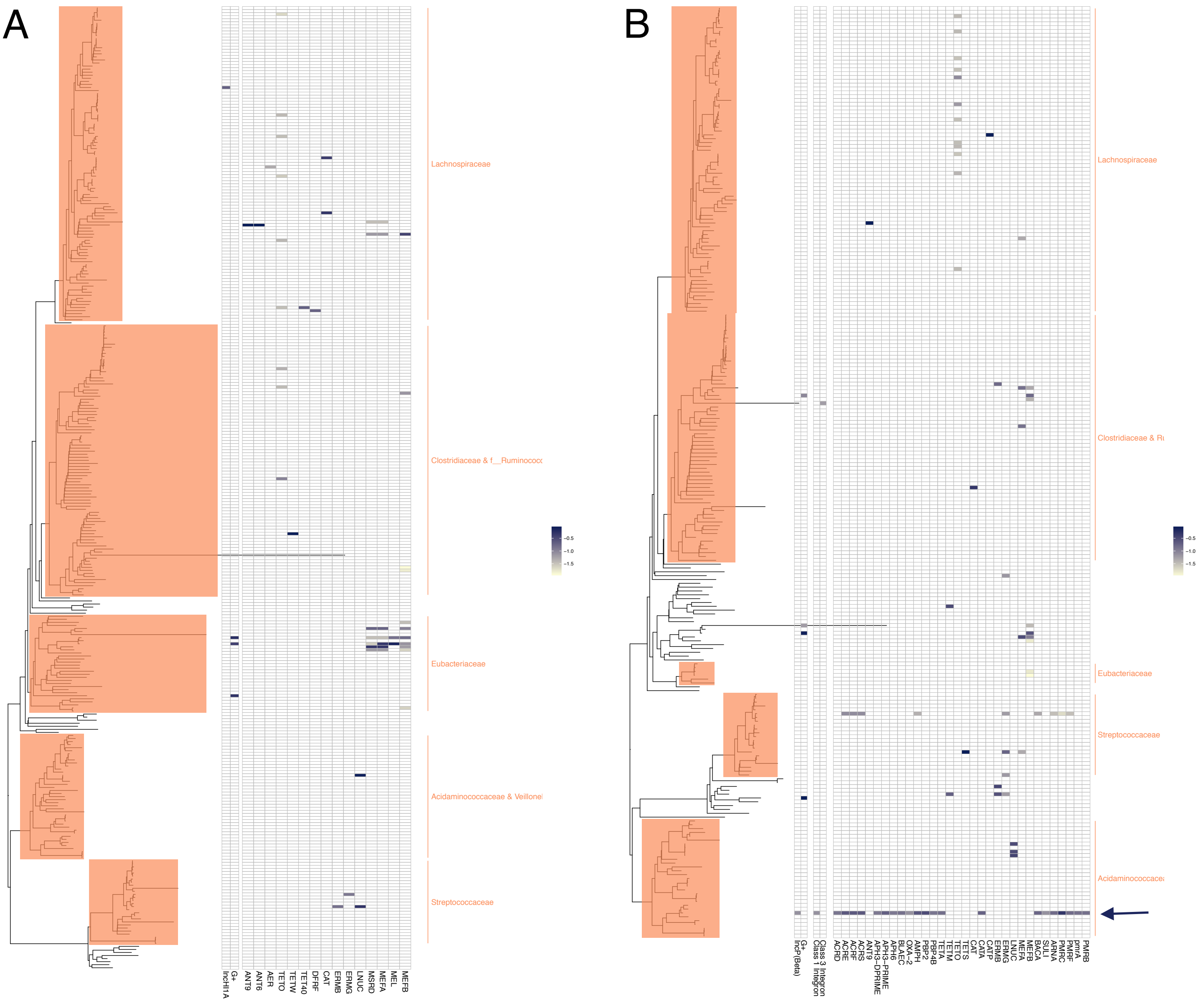
A
B
Lachnospiraceae
Clostridiaceae & f__Ruminococc
Eubacteriaceae
Acidaminococcaceae & Veillonel
Streptococcaceae
IncHI1A
G+
ANT9
ANT6
AER
TETO
TETW
TET40
DFRF
CAT
ERMB
ERMG
LNUC
MSRD
MEFA
MEL
MEFB
-0.5
-1.0
-1.5
Clostridiaceae & Ru
Acidaminococcace
IncP(Beta)
Class 3 Integron
Class 1 Integron
ACRD
ACRE
ACRF
ACRS
ANT9
APH3-PRIME
APH3-DPRIME
APH6
BLAEC
OXA-2
AMPH
PBP2
PBP4B
TETA
TETM
TETO
TETS
CAT
CATA
CATP
ERMB
ERMG
LNUC
MEFA
MEFB
BACA
SULI
ARNA
PMRC
PMRF
pmrA
PMRB
